# After the games are over: life-history trade-offs drive dispersal attenuation following range expansion

**DOI:** 10.1101/014852

**Authors:** T. Alex Perkins, Carl Boettiger, Benjamin L. Phillips

**Affiliations:** Department of Biological Sciences, University of Notre Dame, Notre Dame, IN, USA; Department of Environmental Science, Policy, & Management, University of California, Berkeley, Berkeley, CA, USA; School of Biosciences, University of Melbourne, Melbourne, Victoria, Australia

**Keywords:** competition, evolution, fitness, range expansion, theory

## Abstract

Increased dispersal propensity often evolves on expanding range edges due to the Olympic Village effect, which involves the fastest and fittest finding themselves together in the same place at the same time, mating, and giving rise to like individuals. But what happens after the range’s leading edge has passed and the games are over? Although empirical studies indicate that dispersal propensity attenuates following range expansion, hypotheses about the mechanisms driving this attenuation have not been clearly articulated or tested. Here we use a simple model of the spatiotemporal dynamics of two phenotypes, one fast and the other slow, to propose that dispersal attenuation beyond pre-expansion levels is only possible in the presence of trade-offs between dispersal and life-history traits. The Olympic Village effect ensures that fast dispersers pre-empt locations far from the range’s previous limits. When trade-offs are absent, this pre-emptive spatial advantage has a lasting impact, with highly dispersive individuals attaining equilibrium frequencies that are strictly higher than their introduction frequencies. When trade-offs are present, dispersal propensity decays rapidly at all locations. Our model’s results about the post-colonization trajectory of dispersal evolution are clear and, in principle, should be observable in field studies. We conclude that empirical observations of post-colonization dispersal attenuation offer a novel way to detect the existence of otherwise elusive trade-offs between dispersal and life-history traits.

## Introduction

When a population spreads across space, several evolutionary forces come into play that should drive the evolution of increased dispersal propensity as the invasion front moves forward (Travis and Dytham, 2002). First, under the Olympic Village effect (Phillips et al., 2008), inhabitants at the farthest reaches of the invasion front tend to be limited to the most capable dispersers (Shine et al., 2011; Benichou et al., 2012), leading to spatially assortative mating by dispersal propensity and the perpetuation of this effect in subsequent generations (Phillips et al., 2010). More recently, this phenomenon has been referred to as “spatial sorting” (Shine et al., 2011), which we adopt henceforth. Second, in a density-regulated context, these highly dispersive phenotypes arriving on the invasion front benefit from a fitness advantage through lowered competition with conspecifics (Phillips et al., 2008). Finally, this fitness advantage of more dispersive types may increase over time, as life-history traits also undergo adaptive evolution in the vanguard population (Perkins et al., 2013).

Despite the fact that these distinct evolutionary forces have only recently been elucidated, there is a rapidly growing body of empirical work showing that dispersal propensity often does increase on invasion fronts. Spreading populations ranging from trees to ants, crickets, beetles, and amphibians have all shown evidence of such increases as their ranges have expanded (Cwynar and MacDonald, 1987; Simmons and Thomas, 2004; Léotard et al., 2009; Alford et al., 2009; Lombaert et al., 2014). These rapid increases in dispersal propensity have broad implications for ecological management (*e.g*., the management of invasive species, or native species shifting under climate change) and even medicine (*e.g*., the growth of tumors, and the formation of biofilms; Orlando et al., 2013; van Ditmarsch et al., 2013). Athough there is growing appreciation for the evolution of increased dispersal propensity in the low-density environments along expanding range edges, much less is known about what happens after the range edge has passed and equilibrium densities have been attained.

The evolutionary trajectory of dispersal after colonization is important to understand for at least three key reasons. First, it gives an indication of the long-term consequences of invasion on evolution. If the evolution of increased dispersal is a transient phenomenon, with reversion to pre-invasion levels following colonization, then its long-term implications are modest. If, however, dispersal tends to be maintained at high levels following establishment, then a persistent cline in dispersal phenotypes is expected to exist across the species’ range, with implications for population dynamics and life-history evolution. In this case, invasion may also be a driver of diversification, with many instances of geographic variation being a product of past invasions (Phillips et al., 2010) rather than local adaptation along an underlying environmental gradient (Kirkpatrick and Barton, 1997). Second, understanding the post-colonization trajectory of dispersal could potentially yield further insight into evolutionary processes occurring on the invasion front. Trade-offs between dispersal and fitness, for example, may alter the evolutionary and spread dynamics of the invasion front (Burton et al., 2010; Orlando et al., 2013). If such a trade-off exists, it may manifest as a rapid attenuation of dispersal propensity behind the invasion front. Third, and related to the first point, the reality is that the vast majority of well-documented examples of spread pertain to events that have more or less concluded (Perkins, 2012, Appendix F). Documenting the spread of an invasive species takes time – time in which the population is filling its new range – and so invasions are often only well-documented as they are reaching their conclusion. Because of this, inferences about evolutionary processes on invasion fronts will often be made by examining populations at numerous times post-colonization; a kind of space-for-time substitution (*e.g*., Phillips et al., 2008). Such inferences depend critically on knowing the extent to which populations sampled post-colonization resemble populations on the invasion front when it originally passed that location.

Here we attempt to provide clarity about the processes that govern the evolution of dispersal from the moment of colonization onwards. It is clear from a limited number of theoretical and empirical studies that populations may evolve attenuated dispersal propensity following colonization (Duckworth and Badyaev, 2007; Burton et al., 2010; Lindström et al., 2013). One prominent example in bluebirds attributed post-colonization dispersal attenuation to trade-offs between dispersal and life-history traits (Duckworth and Badyaev, 2007). This is a straightforward invocation of natural selection — less dispersive individuals, who also happen to be less aggressive, invest more in parental care and so increase in frequency over time. Another recent example in cane toads attributed post-colonization dispersal attenuation to spatial sorting (Lindström et al., 2013). This mechanism, whereby dispersal phenotypes are sorted along the strong density cline on the invasion front (Shine et al., 2011; Benichou et al., 2012), posits that the constantly shifting density cline creates a situation whereby the flow of dispersing individuals is asymmetric at any point along the cline. The idea is that this then results in less dispersive individuals from high-density areas outnumbering more dispersive individuals from low-density areas. Although the authors of previous studies of post-colonization dispersal attenuation likely had good reason to make one inference or another, they left the question of the generality of these mechanisms unresolved. Below, we develop a simple model and use it to determine the conditions under which these two mechanisms might operate following colonization. This theoretical analysis not only clarifies the likely importance of these mechanisms, but also provides suggestions about empirical signatures of trade-offs between dispersal and life-history traits that manifest in spatially extended populations.

## Methods

### General ecological model

We consider a species with two phenotypes: slow dispersers with density *N_s_*(*t,x*)at time *t* and location *x*, and fast dispersers with density *N_f_*(*t, x*). For convenience, we drop the notation indicating the specification of these variables over time and space. We assume that each type disperses according to a diffusion process with mean squared displacement per time *D_s_* and *D_f_*, respectively. At any particular location *x* in the absence of immigration and emigration, we model the dynamics of the two types using a Lotka-Volterra competition model with logistic population growth. One departure from this model that we make is that we decompose the intrinsic growth rates, *r_s_* and *r_f_*, as *r_s_* = (*ρ_s_* + *b_s_*) — (*ρ_s_* — *d_s_*) and *r_f_* = (*ρ_f_* + *b_f_*) — (*ρ_f_* — *d_f_*). We interpret *ρ_s_* and *ρ_f_* as birth and death rates of each type when either is at its non-zero equilibrium, *b_s_* and *b_f_* as boosts in birth rates at low density, and *d_s_* and *d_f_* as reductions in death rates at low density (defined on [0,*ρ_s_*] and [0,*ρ_f_*], respectively). Together, these assumptions combine to yield

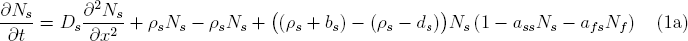

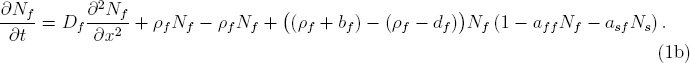

As specified, allowing for non-zero values of *ρ_s_* and *ρ_f_* is inconsequential to the model’s dynamics, but we include them because allowing for turnover within the population at equilibrium is necessary for the model to capture evolutionary change at high density. We now consider one special case and one elaboration of this model that differ in their assumptions about the genetics of the fast and slow phenotypes.

### Genetic models

#### Case 1: Complete heritability

The simplest assumption that can be made about the genetics of these two types is that parents always beget like offspring. One example of when this might be a reasonable approximation would be for a clonal species with a single gene differentiating fast and slow types and infrequent mutation from one type to another relative to the timescale of spatial expansion. When this assumption holds, there is no need to consider non-zero *ρ_s_* or *ρ_f_*, and we can collapse the birth and death rates down to their sums in *r_s_* and *r_f_*. Next, to reduce the number of parameters and to clarify the scales of interest for each variable, we nondimensionalize the model (see Petersen and Hastings (2001) for an overview of this technique in ecology). To do so, we first separate the dimensional and nondimensional components of each variable as 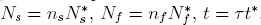, and *x* = *χx*^*^, and then define their dimensional components as 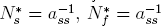, and 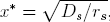, to obtain

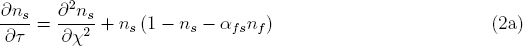

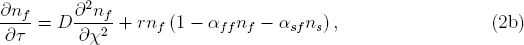

where *D* = *D_f_/D_s_*, *r* = *r_f_/r_s_, αf_s_* = *a_fs_/a_ss_, α_ff_ = a_ff_/a_ss_*, and *α_sf_* = *a_sf_/a_ss_*.

#### Case 2: Single locus in a sexual, diploid species

To consider the possibility that not all offspring will resemble their parental phenotypes, we extend the model to allow for the phenotype to be determined by a pair of alleles at a single locus. We assume that there are only two alleles segregating at this locus, with one resulting in the slow phenotype in individuals homozygous for that allele (the frequency of which is *p*) and the other resulting in the fast phenotype in individuals homozygous for that allele (the frequency of which is *q*). We assume that each heterozygote acquires the slow phenotype with probability *h*. Consequently, there are four states that we must follow: densities of each of the homozygotes, *N_s,pp_* and *N_f,qq_*, and of heterozygotes with either of the phenotypes, *N_s,pq_* and *N_f,pq_*. We assume that differences in birth rates are solely attributable to differences in female fecundity, that mating is random, and that inheritance follows Mendelian proportions.

To follow the dynamics of the four states of interest consistent with these assumptions, we expand on eqns. (1a) and (1b) by taking the following steps: (1) separating *N_s_* into *N_s,pp_* and *N_s,pq_* and *N_f_* into *N_f,pq_* and *N_f,qq_*; (2) adding terms for births of each type owing to matings between parents of all combinations of types in proportions delineated in Table 1; (3) separating dimensional and nondimensional components of each variable; (4) defining dimensional components as 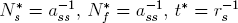, and 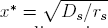, where *r_s_* = *b_s_* + *d_s_*; and (5) performing the necessary algebra to obtain nondimensional equations

**Table 1.** Proportions of matings that result in offspring of a given phenotype and genotype.

| Dam | Sire | Offspring |  |  |  | Dam | Sire | Offspring |  |  |  |
| --- | --- | --- | --- | --- | --- | --- | --- | --- | --- | --- | --- |
| | | $s, pp$ | $s, pq$ | $f, pq$ | $f, qq$ | | | $s, pp$ | $s, pq$ | $f, pq$ | $f, qq$ |
| $\vec{n}_{D,s}$ | $\vec{\eta}_{S,\cdot}$ | $\vec{m}_{s,s,pp}$ | $\vec{m}_{s,s,pq}$ | $\vec{m}_{s,f,pq}$ | $\vec{m}_{s,f,qq}$ | $\vec{n}_{D,f}$ | $\vec{\eta}_{S,\cdot}$ | $\vec{m}_{f,s,pp}$ | $\vec{m}_{f,s,pq}$ | $\vec{m}_{f,f,pq}$ | $\vec{m}_{f,f,qq}$ |
| $n_{s,pp}$ | $\eta_{s,pp}$ | 1 | 0 | 0 | 0 | $n_{f,pq}$ | $\eta_{s,pp}$ | $\frac{1}{2}$ | $\frac{1}{2}h$ | $\frac{1}{2}(1-h)$ | 0 |
| $n_{s,pp}$ | $\eta_{s,pq}$ | $\frac{1}{2}$ | $\frac{1}{2}h$ | $\frac{1}{2}(1-h)$ | 0 | $n_{f,pq}$ | $\eta_{s,pq}$ | $\frac{1}{4}$ | $\frac{1}{2}h$ | $\frac{1}{2}(1-h)$ | $\frac{1}{4}$ |
| $n_{s,pp}$ | $\eta_{f,pq}$ | $\frac{1}{2}$ | $\frac{1}{2}h$ | $\frac{1}{2}(1-h)$ | 0 | $n_{f,pq}$ | $\eta_{f,pq}$ | $\frac{1}{4}$ | $\frac{1}{2}h$ | $\frac{1}{2}(1-h)$ | $\frac{1}{4}$ |
| $n_{s,pp}$ | $\eta_{f,qq}$ | 0 | $h$ | $1-h$ | 0 | $n_{f,pq}$ | $\eta_{f,qq}$ | 0 | $\frac{1}{2}h$ | $\frac{1}{2}(1-h)$ | $\frac{1}{2}$ |
| $n_{s,pq}$ | $\eta_{s,pp}$ | $\frac{1}{2}$ | $\frac{1}{2}h$ | $\frac{1}{2}(1-h)$ | 0 | $n_{f,qq}$ | $\eta_{s,pp}$ | 0 | $h$ | $1-h$ | 0 |
| $n_{s,pq}$ | $\eta_{s,pq}$ | $\frac{1}{4}$ | $\frac{1}{2}h$ | $\frac{1}{2}(1-h)$ | $\frac{1}{4}$ | $n_{f,qq}$ | $\eta_{s,pq}$ | 0 | $\frac{1}{2}h$ | $\frac{1}{2}(1-h)$ | $\frac{1}{2}$ |
| $n_{s,pq}$ | $\eta_{f,pq}$ | $\frac{1}{4}$ | $\frac{1}{2}h$ | $\frac{1}{2}(1-h)$ | $\frac{1}{4}$ | $n_{f,qq}$ | $\eta_{f,pq}$ | 0 | $\frac{1}{2}h$ | $\frac{1}{2}(1-h)$ | $\frac{1}{2}$ |
| $n_{s,pq}$ | $\eta_{f,qq}$ | 0 | $\frac{1}{2}h$ | $\frac{1}{2}(1-h)$ | $\frac{1}{2}$ | $n_{f,qq}$ | $\eta_{f,qq}$ | 0 | 0 | 0 | 1 |

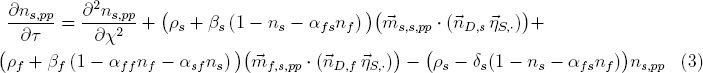

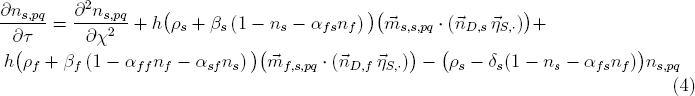

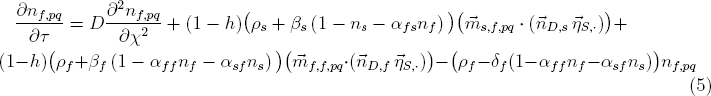

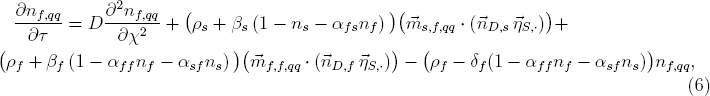

where *ρ_s_* = *ρ_s_/r_s_, ρ_f_* = *ρ_f_/r_s_, β_s_* = *b_s_/r_s_, β_f_* = *b_f_/r_s_, δ_s_* = *d_s_/r_s_, δ_f_* = *d_f_/r_s_, α_fs_* = *a_fs_/a_ss_*, *α_sf_* = *a_sf_/a_ss_*, and *α_ff_* = *a_ff_/a_ss_*, and *η_s,pp_, η_s,pq_, η_f,pq_*, and *η_f,qq_* are normalized frequencies of each type. Vectors of the densities of females of each phenotype-genotype combination 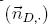, the frequencies of males of each such combination 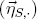, and the proportion of each type of mating resulting in offspring of a given combination 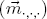 are provided in Table 1.

#### Model analyses

Our primary interest was understanding patterns of the relative frequencies of the fast and slow types across space long after initial colonization. This required first determining the range of patterns that are possible and then assessing how different ecological scenarios affect those patterns. Because our interests were general and not in reference to any particular system, we limited our analyses to the nondimensionalized equations, which emphasize relative differences between the two types.

We solved each of the models under scenarios in which there was either no trade-off (*i.e*., all life-history parameters equal for both types) or a trade-off in any of four life-history parameters (*i.e*., differences in birth, death, or competition resulting in lower population growth of the fast type), implemented one at a time with the following values: *r* = 0.8 (or *β_f_* = *δ_f_* = 0.4), *α_ff_* = 1.1, *α_fs_* = 0.9, and *α_sf_* = 1.1. It is not clear a priori whether certain types of trade-offs should impact patterns of post-colonization dispersal evolution in different ways, and so we considered all possible trade-offs under our simple ecological model. Unless specified otherwise, *D* = 1.2 was used as a default relative difference in dispersal, meaning that fast individuals have a 20% higher dispersal coefficient than slow ones. Whenever a life-history trade-off was not being implemented, default values of life-history parameters were *r* = 1, *β_s_* = *β_f_* = *δ_s_* = *δ_f_* = 0.5, *α_fs_* = *α_sf_* = *α_ff_* = 1, and *ρ_s_* = *ρ_f_* = 1 or 0. In addition, we analyzed each scenario about life-history trade-offs under different assumptions about density dependence. For the model in eqns. (2a) & (2b), the possibilities were either density-independent or density-dependent growth. For the model in eqns. (3)-(6), the possibilities included either density-independent growth or density-dependent growth with *ρ_s_* = *ρ_f_* = 0 or 1. We limited analyses of the latter model to *h* = 0.5.

We solved the model in eqns. (2a) & (2b) on a space-time domain of [0, 40]_χ_ x [0,100]_τ_ for density-independent growth and [0, 40]_χ_ x [0, 400]_τ_ for density-dependent growth, both under initial conditions of *n_s_* = *n_f_* = 0.1 at *χ* = 0 and *n_s_* = *n_f_* = 0 elsewhere. Conditions for the model in eqns. (3)-(6) were the same but with the additional specification that all η = 0.25 at τ = 0, which is consistent with *n_s_* = *n_f_* and *h* = 0.5 in a population at Hardy-Weinberg equilibrium. All solutions of the models were obtained numerically using the deSolve package (Soetaert et al., 2010) in R (R Core Team, 2014).

## Results

### Aspatial model: equilibrium properties

Before we examine dynamics under the spatial models, we first note the equilibrium properties of the nondimensionalized ecological model in a population with no emigration or immigration, as this is useful background information for interpreting the spatial results. Under default parameter values in which there are no life-history trade-offs, there are infinitely many unstable equilibria 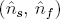 satisfying 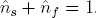. In the more general case where some or all *α* ≠ 1, the equilibrium values of the two types in an isolated local population are either 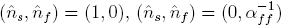, or, in cases where the parameter values yield positive values for both types,

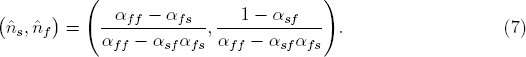

Under this latter case, in which some or all *α* ≠ 1, the equilibrium and stability properties of the model are equivalent to those of the Lotka-Volterra model. The special case in which all *α* = 1 is not one that has been emphasized in studies of interspecific competition, but it is highly appropriate for examining the dynamics of two or more types of a single species with similar, but potentially differing, life-history properties.

### Spatial model: density-independent growth

Under either model with density-independent population growth, the frequency of the slow type always approaches an equilibrium that is constant across the entire spatial domain (Figs. 1 & 2). When the intrinsic growth rates of the two types are equal (indicated by the ratio r = 1), the initial frequencies are approached across the entire domain as time *τ* → ∞ (Figs. 1 & 2, top left). In the density-independent case, there is a persistent density gradient present, even as *τ* → ∞. This density gradient sets up conditions for a net flux of genotypes from high-density to low-density areas; thus, we see the slow type invade and the long-term frequencies slowly approach the initial frequencies across the entire domain.

**Figure 1.**
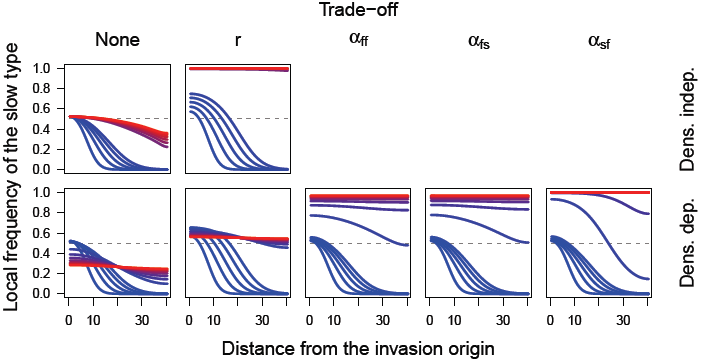
Spatial spread of the slow type under different scenarios about density dependence and life-history trade-offs under the model in eqns. (2a) & (2b). Each curve shows the invasion profile of the slow type at a given point in time, with time indicated by colors ranging from blue at time τ = 0 to red at time τ = 100 in the top row and τ = 400 in the bottom row.

**Figure 2.**
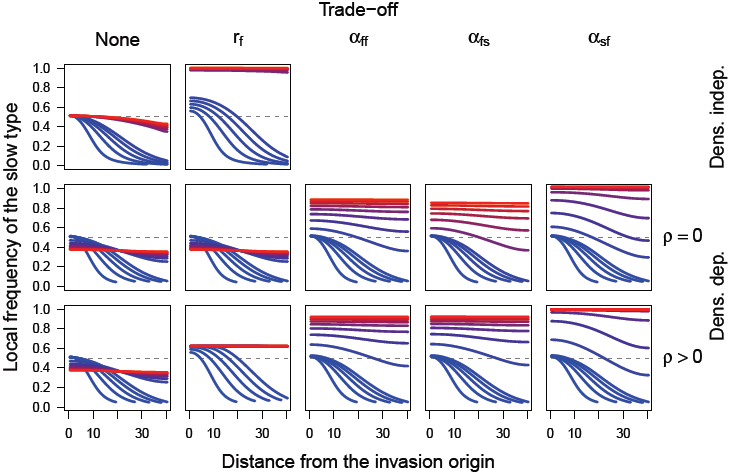
Spatial spread of the slow type under different scenarios about density dependence and life-history trade-offs under the model in eqns. (3)–(6). Each curve shows the invasion profile of the slow type at a given point in time, with time indicated by colors ranging from blue at time τ = 0 to red at time τ = 100 in the top row and τ = 400 in the bottom two rows.

### Spatial model: density-dependent growth

The frequency of the slow type always approaches a constant equilibrium under models with density dependence, as well (Figs. 1 & 2). In this case, when the intrinsic population growth rates of the two types are equal (*i.e*., *β_s_* = *β_f_* = *δ_s_* = *δ_f_* = 0.5), the equilibrium frequency of the slow type is always less than its frequency at introduction (Figs. 1 & 2, bottom left). The reason that this equilibrium frequency is always less than the introduction frequency, unlike under the density-independent model, is that growth ceases once equilibrium densities are attained. Thus, any numerical advantages that the fast type enjoys at locations far from the invasion origin are preserved in the long term, because any 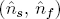 satisfying 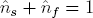 is an equilibrium. However, because those equilibria are unstable, they shift in response to perturbations from dispersal from nearby locations with slightly different frequencies, eventually resulting in a homogenization of frequencies across space. The value of this spatially homogenized equilibrium frequency of the slow type depends on the initial frequency of the slow type, the relative dispersal advantage of the fast type, and the extent of the spatial domain (Fig. 3). With a greater relative dispersal ability and a larger spatial domain, the fast type will enjoy preemption of a greater area for a longer time, resulting in a lower equilibrium frequency of the slow type (Fig. 3).

**Figure 3.**
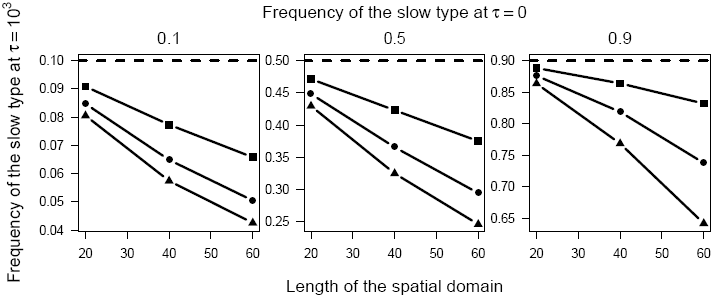
Frequency of the slow type across the entire spatial domain long after all locations have been invaded and equilibrium densities at each location have been attained (*i.e*., τ = 10^3^). These frequencies, which correspond to the frequency indicated by the red line in the bottom left panel of Figure 2, are shown here as a function of the length of the spatial domain (x-axis), the initial frequency of the slow type (dashed lines, separate panels), and the relative dispersal ability of the fast type (square: *D* = 1.1, circle: *D* = 1.2, triangle: *D* = 1.3).

Under all scenarios that we examined, life-history trade-offs always resulted in an increase in the equilibrium frequency of the slow type relative to what it was in the absence of the trade-off (Figs. 1 & 2). For a trade-off in the intrinsic growth rate (indicated by the ratio of growth rates of fast and slow types *r* < 1), the boost in the equilibrium frequency of the slow type attributable to this trade-off is modest, because the trade-off ceases to operate once equilibrium densities are attained (Fig. 1 & 2, second columns, bottom rows). Much like the intermediate equilibrium frequency of the slow type in the absence of trade-offs (Fig. 3), the value of this equilibrium frequency is likely subject to the initial frequency of the slow type, the relative dispersal ability of the fast type, and the extent of the spatial domain, as well as the strength of the trade-off. Other trade-offs that we examined had much clearer effects, leading to eventual fixation of the slow type (Figs. 1 & 2, three rightmost columns). Unlike a trade-off in the intrinsic growth rate, the effect of trade-offs in three different forms of relative competitive ability persisted after overall population growth slowed. The long-term outcome of fixation of the slow type is therefore inevitable based on the equilibrium properties of the competition model given values of the interaction coefficients associated with trade-offs between life-history and dispersal traits.

## Discussion

When populations spread into new areas, more dispersive individuals will often dominate the invasion front and lead to increases in dispersal propensity at the front as it advances (Travis and Dytham, 2002; Phillips et al., 2010). But what happens to dispersal traits post-colonization; *i.e*., after the front has passed? If dispersal propensity attenuates back to pre-invasion levels, why does this happen? Our model captures the two possible drivers of dispersal evolution in this scenario, spatial sorting and natural selection due to life-history trade-offs, and clarifies when and how they operate. A lack of clarity about the operation of these two drivers has impeded the interpretation of empirical results, which have typically assumed one mechanism over the other without fully considering both. A pattern of declining dispersal post-colonization in bluebirds, for example, was attributed entirely to life-history trade-offs (Duckworth and Badyaev, 2007), whereas a similar pattern in cane toads was attributed to spatial sorting (Lindström et al., 2013). Rather than leave this determination to the proclivities of a study’s authors, our goal is to establish a general understanding of how the processes of spatial sorting and natural selection due to life-history trade-offs give rise to these patterns in different contexts.

Our results indicate that both hypotheses about dispersal attenuation behind invasion fronts appear to be at play, but under different circumstances. Spatial sorting operates when there is a persistent density gradient on the invasion front. When we maintained this gradient in the model by allowing the population to grow in a density-independent manner, the frequency of slow dispersers increased after colonization, even in the absence of natural selection (*i.e*., no trade-off between dispersal and fitness; Figs 1 & 2, upper left). Thus, in the early stages of colonization when population growth is exponential, spatial sorting could be an important mechanism driving dispersal attenuation. In the density-dependent case, dispersal also attenuated strongly when a life-history trade-off operated at high densities and when traits were being observed long after the initial colonization of a location. Under those circumstances, life-history trade-offs have plenty of time to operate, the mark of spatial sorting has long vanished (after many generations of gene flow in the absence of a gradient in population density), and a spatially homogenous, slow phenotype will dominate. By contrast, when life-history trade-offs only operate at low densities or are absent altogether, the long-term outcome of dispersal evolution is more complicated. In this case, we observed the spread of fast phenotypes from relatively recently invaded areas back towards the invasion origin and vice versa (Figs 1 & 2, lower left). This happens because, in the absence of a density gradient or a fitness trade-off with dispersal propensity, dispersal acts solely as a homogenizing force across the range.

Quantitative details about the long-term frequencies of dispersal types will depend on additional subtleties, including how quickly equilibrium densities are attained relative to the timescale of spatial spread. Our results show that long-term, range-wide frequencies of fast and slow types are potentially quite sensitive to the relative dispersal advantage of the fast type and the extent of the spatial domain being invaded. With empirical estimates indicating that spread often proceeds for on the order of 10–100 generations (Perkins, 2012, Appendix F) and with mounting evidence of extensive variability in dispersal traits (Hughes et al., 2003; Simmons and Thomas, 2004; Léotard et al., 2009; Lindström et al., 2013; Lombaert et al., 2014), there is good reason to suspect that spatial sorting should guarantee fast types a lasting advantage over their slow counterparts in many instances of range expansion in nature. After all, we have shown (in the density-independent case) that, at best, spatial sorting alone can only restore dispersal propensity back to pre-invasion levels. Spatial sorting alone cannot account for gains by the slow type beyond pre-invasion levels, and only then in the unlikely case of a complete lack of density regulation.

Together, our results suggest that the most likely explanation for empirically observed declines in dispersal following invasion is natural selection due to trade-offs between dispersal and life-history traits. In the bluebird example, aspects of that species’ natural history consistent with this explanation include that local populations are strongly density-regulated (by limited nest sites), the process of re-colonization by the slow type happens within only a few generations, and there is a strong genetic correlation between dispersal propensity and life-history traits, particularly at high densities (Duckworth and Badyaev, 2007; Duckworth, 2008; Duckworth and Kruuk, 2009). Aspects of the cane toad’s natural history are also consistent with our conclusion about the necessity of trade-offs for the evolution of dispersal attenuation. Extensive variation in the dispersal propensities of cane toads, a well-established genetic basis of variation in dispersal and life-history traits, and a spread phase that has unfolded over vast distances and dozens of generations (Phillips et al., 2008; Alford et al., 2009; Lindström et al., 2013) are all conditions that, in the absence of life-history trade-offs, appear highly unfavorable for post-colonization dispersal attenuation. Based on our results, the most likely mechanism for the observation of post-colonization dispersal attenuation in cane toads is the presence of trade-offs between dispersal and fitness. This expectation of a trade-off is supported by recent observations that highly dispersive invasion-front toads have a lower reproductive rate than their conspecifics from the range core (Hudson et al., 2015). It is still possible, however, that spatial sorting was an important driver of post-colonization dispersal attenuation in cane toads, and we note that additional empirical observations guided by our theoretical predictions could help resolve this question in the future (Table 2).

**Table 2.** Interpreting empirical observations in light of theoretical results described here.

| Empirical observation | Theoretical interpretation |
| --- | --- |
| Before equilibrium density has been attained at a site post-colonization, modest dispersal attenuation not beyond pre-invasion levels. | Spatial sorting is definitely operating, and a trade-off between dispersal propensity and fitness may or may not be operating. |
| Before equilibrium density has been attained at a site post-colonization, rapid dispersal attenuation beyond pre-invasion levels. | A trade-off between dispersal propensity and fitness is definitely operating, and spatial sorting may or may not be operating. |
| After equilibrium densities have been attained range-wide, attenuating dispersal beyond pre-invasion levels across the range. | A trade-off between dispersal propensity and fitness is definitely operating, and spatial sorting is definitely not operating. |
| After equilibrium densities have been attained range-wide, increasing dispersal near the invasion origin and attenuating dispersal in more recently invaded populations. | A trade-off between dispersal propensity and fitness is not operating. Instead, gene flow is homogenizing dispersal phenotypes across the range. |

Evaluating the extent to which different conditions apply in recently expanded, spatially distributed species, such as the aforementioned bluebirds and cane toads, suggests a tantalizing possibility: that patterns of dispersal evolution following range expansion could be used to infer the existence of trade-offs between dispersal and life-history traits. Such trade-offs are often posited in theoretical studies (Ronce, 2007), but empirical evidence of their existence is scarce, coming primarily from flight-fecundity trade-offs in insects (*e.g*., Hughes et al., 2003; Duthie et al., 2015). A primary reason for this paucity of examples is that it is logistically difficult to measure relevant variables (life-history and dispersal traits) and then to be sure that all relevant life-history traits have been taken into account (Ronce, 2007; Phillips et al., 2010). Negative relationships between fecundity and dispersal, for example, could be cancelled out by negative correlations between fecundity and age to maturity, which would go undetected unless all traits are measured. Thus, observation of post-colonization dispersal attenuation could provide a novel, and very useful, clue about the existence of trade-offs, even if the proximate traits remain unidentified.

There are, however, a number of limitations that must be kept in mind when reconciling empirical results with those from our theoretical study. Perhaps the most important limitation is that it could take a very long time for the long-term behavior that we studied to supplant prolonged periods of transient behavior. In particular, our model ultimately does not allow for stable clines in dispersal propensity, yet clines are clearly manifest across many species’ ranges for years after colonization (Cwynar and MacDonald, 1987; Simmons and Thomas, 2004; Léotard et al., 2009; Alford et al., 2009; Lombaert et al., 2014). Temporal trends in dispersal clines measured at fixed locations across a recently established range should nonetheless yield empirical signatures consistent with one model scenario or another. There is also a tremendous amount of detail that our model eschews, including demographic stochasticity, mutation, mating system, relatedness, and Allee effects, all of which have known relevance to dispersal evolution (Cadet et al., 2003; Burton et al., 2010; Travis et al., 2010; Hargreaves and Eckert, 2013; Shaw and Kokko, 2015). Such details will undoubtedly play an important role in future studies that use system-specific models to evaluate the quantitative plausibility of alternative hypotheses about the drivers and consequences of dispersal evolution in a given natural system (*e.g*., Perkins et al., 2013).

Although system-specific modeling and observation will be crucial for future studies, our analysis provides an important first suggestsion that dispersal propensity typically only shows long-term attenuation in density-regulated populations when there is a trade-off between dispersal and fitness operating at high density. If these trade-offs are prevalent in nature, then the long-term implications of dispersal evolution during invasion are likely modest. Gradients in dispersal across the invaded range will, in the absence of alternative fitness peaks, ultimately be transient phenomena, and diversification of life-histories driven by spread may be unusual. Thus, the existence of trade-offs is critical to determine. Importantly, our model suggests that dispersal phenotypes at a location will rarely reflect the phenotypes that first colonized that location, and the course of evolution after colonization can be used to determine whether or not a trade-off is operating. Although we do not yet know how prevalent trade-offs between dispersal and life-history traits are in nature, we now have a new way of looking for them.

## References

Alford, R., G. Brown, L. Schwarzkopf, B. Phillips, and R. Shine. 2009. Comparisons through time and space suggest rapid evolution of dispersal behaviour in an invasive species. Wildlife Research 36:23–28.

Benichou, O., V. Calvez, N. Meunier, and R. Voituriez. 2012. Front acceleration by dynamic selection in Fisher population waves. Physical Review E 86:041908.

Burton, O. J., J. M. J. Travis, and B. L. Phillips. 2010. Trade-offs and the evolution of life-histories during range expansion. Ecology Letters 13:1210–1220.

Cadet, C., R. Ferriere, J. A. J. Metz, and M. van Baalen. 2003. The evolution of dispersal under demographic stochasticity. American Naturalist 162:427–442.

Cwynar, L. L. C., and G. M. G. MacDonald. 1987. Geographical variation of lodgepole pine in relation to population history. American Naturalist 129:463–469.

Duckworth, R. A. 2008. Adaptive dispersal strategies and the dynamics of range expansion. American Naturalist 172:S4–S17.

Duckworth, R. A., and A. V. Badyaev. 2007. Coupling of dispersal and aggression facilitates the rapid range expansion of a passerine bird. Proceedings of the National Academy of Sciences 104:15017–22.

Duckworth, R. A., and L. E. B. Kruuk. 2009. Evolution of genetic integration between dispersal and colonization ability of a bird. Evolution 63:968–977.

Duthie, A. B., K. C. Abbott, and J. D. Nason. 2015. Trade-offs and coexistence in fluctuating environments: evidence for a key dispersal-fecundity trade-off in five nonpollinating fig wasps. American Naturalist 186:151–158.

Hargreaves, A. L., and C. G. Eckert. 2013. Evolution of dispersal and mating systems along geographic gradients: implications for shifting ranges. Functional Ecology 28:5–21.

Hudson, C. M., B. L. Phillips, G. P. Brown, and R. Shine. 2015. Virgins in the vanguard: low reproductive frequency in invasion front toads. Biological Journal of the Linnean Society in press.

Hughes, C. L., C. Dytham, and J. K. Hill. 2003. Evolutionary trade-offs between reproduction and dispersal in populations at expanding range boundaries. Proceedings of the Royal Society Biological Sciences Series B 270:S147–S150.

Kirkpatrick, M., and N. H. Barton. 1997. Evolution of a species’ range. American Naturalist 150:1–23.

Léotard, G., G. Debout, A. Dalecky, S. Guillot, L. Gaume, D. McKey, and F. Kjellberg. 2009. Range expansion drives dispersal evolution in an equatorial three-species symbiosis. PLOS One 4:e5377.

Lindström, T., G. P. Brown, S. A. Sisson, B. L. Phillips, and R. Shine. 2013. Rapid shifts in dispersal behavior on an expanding range edge. Proceedings of the National Academy of Sciences 110:13452–13456.

Lombaert, E., A. Estoup, B. Facon, B. Joubard, J.-C. Grégoire, A. Jannin, A. Blin, and T. Guillemaud. 2014. Rapid increase in dispersal during range expansion in the invasive ladybird Harmonia axyridis. Journal of Evolutionary Biology 27:508–517.

Orlando, P. A., R. A. Gatenby, and J. S. Brown. 2013. Tumor evolution in space: the effects of competition colonization tradeoffs on tumor invasion dynamics. Frontiers in Oncology 3:1.

Perkins, T. A. 2012. Evolutionarily labile species interactions and spatial spread of invasive species. American Naturalist 179:E37–E54.

Perkins, T. A., B. L. Phillips, M. L. Baskett, and A. Hastings. 2013. Evolution of dispersal and life-history interact to drive accelerating spread of an invasive species. Ecology Letters 16:1079–1087.

Petersen, J. E., and A. Hastings. 2001. Dimensional approaches to scaling experimental ecosystems: designing mousetraps to catch elephants. American Naturalist 157:324–333.

Phillips, B., G. Brown, and R. Shine. 2010. Life-history evolution in range-shifting populations. Ecology 91:1617–1627.

Phillips, B. L., G. P. Brown, J. M. J. Travis, and R. Shine. 2008. Reid’s paradox revisited: the evolution of dispersal kernels during range expansion. American Naturalist 172 Suppl:S34–48.

R Core Team, 2014. R: A Language and Environment for Statistical Computing. R Foundation for Statistical Computing, Vienna, Austria.

Ronce, O. 2007. How does it feel to be like a rolling stone? Ten questions about dispersal evolution. Annual Review of Ecology, Evolution, and Systematics 38:231–253.

Shaw, A. K., and H. Kokko. 2015. Dispersal evolution in the presence of allee effects can speed up or slow down invasions. American Naturalist 185:631–639.

Shine, R., G. P. Brown, and B. L. Phillips. 2011. An evolutionary process that assembles phenotypes through space rather than through time. Proceedings of the National Academy of Sciences 108:5708–5711.

Simmons, A. D., and C. D. Thomas. 2004. Changes in dispersal during species’ range expansions. American Naturalist 164:378–395.

Soetaert, K., T. Petzoldt, and R. W. Setzer. 2010. Solving differential equations in r: Package desolve. Journal of Statistical Software 33:1–25.

Travis, J. M. J., and C. Dytham. 2002. Dispersal evolution during invasions. Evolutionary Ecology Research 4:1119–1129.

Travis, J. M. J., T. Münkemüller, and O. J. Burton. 2010. Mutation surfing and the evolution of dispersal during range expansions. Journal of Evolutionary Biology 23:2656–2667.

van Ditmarsch, D., K. E. Boyle, H. Sakhtah, J. E. Oyler, C. D. Nadell, E. Déziel, L. E. Dietrich, and J. B. Xavier. 2013. Convergent evolution of hyperswarming leads to impaired biofilm formation in pathogenic bacteria. Cell Reports 4:697–708.

